# The role of alpha oscillations in a premotor-cerebellar loop in modulation of motor learning: insights from transcranial alternating current stimulation

**DOI:** 10.1101/2020.07.27.209148

**Authors:** Christine Schubert, Alhuda Dabbagh, Joseph Classen, Ulrike M. Krämer, Elinor Tzvi

## Abstract

Alpha oscillations (8-13 Hz) have been shown to play an important role in dynamic neural processes underlying learning and memory. The goal of this study was to scrutinize the role of α oscillations in communication within a network implicated in motor sequence learning. To this end, we conducted two experiments using the serial reaction time task. In the first experiment, we explored changes in α power and cross-channel α coherence. We found a gradual decrease in learning-related α power over left premotor cortex (PMC), somatosensory cortex (S1) and tempo-parietal junction (TPJ). Alpha coherence between left PMC/S1 and right cerebellar crus I was reduced for sequence learning, possibly reflecting a functional decoupling in a motor-cerebellar loop during the motor learning process. In the second experiment in a different cohort, we applied 10Hz transcranial alternating current stimulation (tACS), a method shown to entrain local oscillatory activity, to left M1 (lM1) and right cerebellum (rCB) during sequence learning. We observed learning deficits during rCB tACS compared to sham, but not during lM1 tACS. In addition, learning-related α power following rCB tACS was increased in left PMC, possibly reflecting a decrease in neural activity. Importantly, learning-specific coherence between left PMC and right cerebellar lobule VIIb was enhanced following rCB tACS. These findings suggest that interactions within a premotor-cerebellar loop, which are underlying motor sequence learning, are mediated by α oscillations. We show that they can be modulated through external entrainment of cerebellar oscillations, which then modulates motor cortical α and interferes with sequence learning.

## Introduction

Neural oscillations have gained increasing interest as a potential key mechanism underlying cognitive functions. Specifically, alpha oscillations (α, 8-13Hz) have been linked to working memory (Bonnefond and Jensen, 2012; Jensen, 2002; Sauseng et al., 2010) as well as long-term memory processes (Meeuwissen et al., 2011). Oscillations in the α frequency range over motor areas, also known as μ oscillations, were shown to decrease with motor sequence learning (Boenstrup et al., 2014) as well as motor memory consolidation (Pollok et al., 2014). During learning of items in a sequence, occipito-parietal α power increase was associated with faster reaction times, suggesting the critical role of oscillatory α in forming top-down sensory predictions of the upcoming stimuli (Crivelli-Decker et al., 2018). Changes in α power were also linked to pre-stimulus preparatory processes over frontal areas during learning, as well as to memory consolidation, in occipito-parietal areas (Moisello et al., 2013). Finally, we previously showed that during visuomotor sequence learning, α power decreases over occipito-parietal areas as learning progresses (Tzvi et al., 2018, 2016). Together, this evidence suggests that for both motor and perceptual sequence learning dynamic changes in α power at functional relevant sites may play a critical role.

To further link α oscillations to motor sequence learning, previous studies have employed transcranial alternating current stimulation (tACS) at 10 Hz (α mid-frequency) and investigated consequent performance changes. Indeed, evidence suggests that 10 Hz tACS leads to local entrainment of oscillatory activity (Zaehle et al., 2010) and modulation of cognitive functions (Helfrich et al., 2014). A recent, comprehensive study in primates showed that while tACS (at different frequencies including 10 Hz) does not affect firing rates of individual cells or cell populations, it does entrain activity in single neurons, specifically for the targeted frequency and spatial location (Krause et al., 2019).

In motor sequence learning, 10Hz tACS to left primary motor cortex (M1) led to enhanced performance in some studies (Antal et al., 2008; Pollok et al., 2015) but disrupted consolidation in others (Rumpf et al., 2019) with conflicting effects depending on age (Fresnoza et al., 2020). Importantly these studies could not establish a link between learning and actual oscillatory entrainment in the α range. Thus, the extent to which α oscillations affect motor learning remains to be established.

In previous fMRI studies (Tzvi et al., 2015, 2014), we showed that interactions in a cortico-cerebellar loop are critical for motor learning. The goal of the current study was to investigate whether α oscillations mediate interactions in this network and thereby enable motor learning. To this end, we performed two EEG experiments in independent samples, using a modified version of the serial reaction time task (SRTT) (Nissen and Bullemer, 1987). In the first experiment, we investigated α power changes during motor sequence learning, as well as α-dependent connectivity using coherence. In the second experiment, we applied 10Hz tACS (i.e., in the midrange of α) to left M1 and right cerebellum during motor sequence learning and examined subsequent oscillatory changes using EEG. We hypothesized that tACS to either M1 or cerebellum would lead to local entrainment of α oscillations and consequently changes in learning-related α power, measured following tACS. In terms of behavior, we expected that these oscillatory changes affect learning performance during and following tACS.

## 1. Materials and Methods

### 1.1. Experiment 1

#### 1.1.1. Participants

Twenty-six healthy participants (mean age: 23 years, range 18-38; 9 males) took part in the experiment after giving informed consent. Participants were financially compensated or received course credit. All participants were right-handed, had normal or corrected to normal vision with no color deficiency. We excluded participants who regularly played a musical instrument or computer games. The study was approved by the Ethics Committee of the University of Lübeck and was performed in accordance with the Declaration of Helsinki. One participant was excluded due to an error in data acquisition. An additional participant was excluded due to high error-rates (see behavioral analysis). For the EEG data analysis, two participants were excluded due to artifacts in the recorded EEG signal, resulting in a sample of 22 participants for the EEG analyses of Experiment 1.

#### 1.1.2. Experimental paradigm and task design

During the EEG recordings and task performance, participants were seated comfortably in front of a 17” screen, about 1 m away, on which visual stimuli were presented. Before and after the experimental blocks, we recorded resting-state sessions, which were not further analyzed for the present work. Participants performed a modified version of the SRTT. In each trial, four squares were presented on a grey background in a horizontal array, with each square (from left to right) associated with the four fingers of the right hand (Fig. 1A). At stimulus onset, one of the squares turned blue and the rest remained black. Subjects were instructed to respond to this blue colored square with the corresponding button, as precisely and quickly as possible. The stimulus remained on the screen until a button press was registered. In case of a wrong button press, the blue colored square turned to red to mark the error. The response-stimulus interval was 500 ms (Fig. 1B). Trials were counted as correct when the appropriate key was pressed within 1000 ms after stimulus onset. In case no button was pressed within this time frame, a text appeared on the screen requesting the participants to be faster (“Schneller!”).

**Figure 1.**
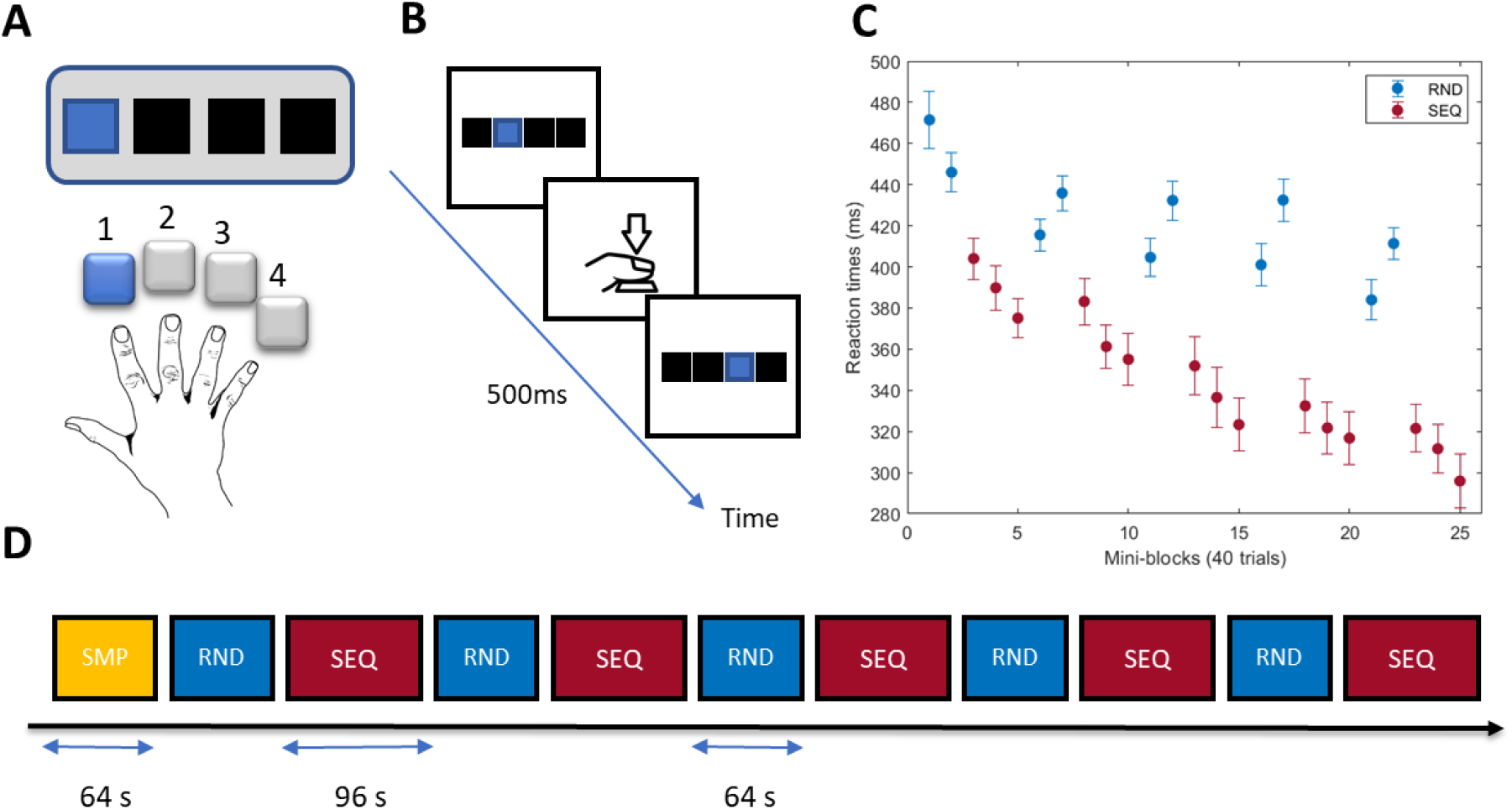
Experiment 1: Experimental design and behavioural results. **A** Serial reaction time task. In each trial, 4 black squares were presented. At stimulus onset, one of the squares turned blue and subjects were instructed to press the button corresponding to the blue square with the respective finger. **B** Task timeline. **C**. Reaction times averaged across subjects in each mini-block with 40 trials, and each condition. Error bars are standard errors of the mean. RND – random trials. SEQ – sequence trials. **D** Subjects performed five RND blocks of 80 trials and five SEQ blocks of 120 trials in an alternating order. A simple block (SMP – yellow) with 80 trials appeared before or after the main task.

The task consisted of three different conditions: simple (SMP), random (RND), and sequence (SEQ). In SMP, stimuli were presented in a simple order of button presses 4-3-2-1-4-3-2-1. In RND, stimuli were presented in pseudorandom order, generated using Matlab (Natick, MA), such that items appeared exactly twice and were not repeated. In SEQ, stimuli were organized in an 8-items-sequence in a second order predictive manner, which means that pairs of consecutive stimuli, rather than single stimuli (which would be first-order predictive) are followed by some other stimuli, thereby preventing learning by pairwise associations (Curran, 1997). The task contained a total number of 11 blocks, separated by 20 s breaks. There was one SMP block at the beginning or the end of the task (counter-balanced between subjects), and five RND blocks and five SEQ blocks in an alternating order (see Fig. 1D for an exemplary order). Each of the SEQ blocks contained 15 repetitions of the 8-element sequence summing to a total of 120 trials per block. The SMP block contained 10 repetitions of the simple sequence summing to a total of 80 trials. Each of the RND blocks contained 80 trials.

Next and after a short (~5 min) break, participants performed a completion task to assess explicit knowledge of the sequence. In this task, the exact same set of stimuli of the SRTT was presented. The 8-element sequence was repeated 32 times. In each repetition one regular trial was substituted by a completion trial. In a completion trial, the target square was replaced by a question mark and subjects had to press a button corresponding to one of the 4 squares which they believed should be the next target square. Each position in the sequence was therefore tested four times producing 32 completion trials. After guessing, subjects were asked whether they were sure of their choice and gave a YES/NO answer. This procedure enabled us to differentiate between a correct response and a correct assured response.

#### 1.1.3. Behavioral analysis

We computed reaction times (RTs) and error-rates (ERs) for each of the experimental conditions. Both wrong button-presses and missing responses were regarded as errors. RTs were averaged across mini-blocks of 40 trials, corresponding to five repetitions of the 8-element sequence in SMP and SEQ. This sub-division resulted in two mini-blocks for each RND and SMP block and three mini-blocks for each SEQ block. We excluded trials in which the participants made an error as well as trials in which RTs were either longer than 1000 ms or deviated by more than 2.7 standard deviations (SD) from their average response time (corresponding to p < 0.01). To measure motor sequence learning in each block, we used the difference between the averaged RT in the second mini-block of each RND block with the first mini-block of the following SEQ block (Fig. 1C). These values were then entered into a 2 x 5 repeated measures ANOVA (rmANOVA) with factors COND (SEQ, RND) and TIME (5). The ERs were computed by dividing the number of errors in each block by the number of trials in that block. As these data were not distributed normally, we tested differences between SEQ and RND in each block using the signrank test. Note that in this analysis there was no sub-division into mini-blocks.

#### 1.1.4. EEG recordings

EEG was recorded with Ag/AgCl electrodes and a 64-channel BrainAmp MR plus amplifier with a sampling rate of 250 Hz. Electrodes were placed according to an extension of the international 10–20 system. Vertical and horizontal eye movements (vEOG and hEOG) were recorded, using electrodes placed below the right eye and a frontopolar electrode (vEOG), as well as electrodes located on the outer canthus of each eye (hEOG). The EEG was recorded against a reference electrode placed on the right earlobe and a ground electrode in FPz. All electrode impedances were kept below 5 kΩ.

#### 1.1.5. EEG pre-processing

Pre-processing and all subsequent analyses were performed using in-house Matlab (The Mathworks®, Natick, MA) scripts and the EEGLAB toolbox. EEG signals were re-referenced offline to the average of the signal from the two earlobe electrodes. A high-order band-pass filter (*Fcutoff* = 1 – 49 Hz) was applied to the signals to remove slow drifts and power line noise. Then, the signals were segmented into 3 s epochs (−1 s to 2 s around stimulus onset). We then identified noisy electrodes especially in locations Fp1, Fp2, T7, T8 and EOG and interpolated them. An independent component analysis (ICA) procedure was then applied to the signals to identify components related to eye blinks and horizontal eye movements based on their topography and signal shape. In most subjects we removed 2-6 components. Additional artifacts were removed using a simple threshold (−80 μV, +80 μV) on the filtered data.

#### 1.1.6. Spectral power and source analysis

Next, we computed the power spectrum of the EEG signals using the Morlet wavelet as implemented in Fieldtrip (http://www.fieldtriptoolbox.org). Data were filtered to obtain oscillatory power between 1 Hz and 50 Hz using wavelets of 7 cycle length. To evaluate the effect of learning on EEG power, we computed power differences between SEQ and RND trials in each consecutive block (time point TP1-TP5), averaged for a pre-stimulus period across a time window of −300 −0 ms (0 ms being stimulus onset), and for post-stimulus period across a time window of 0 – 300 ms. Note that these time windows were selected to minimize overlap with button presses. Statistical tests were performed by subjecting power differences (SEQ-RND) in TP1-TP5 to a 1-way rmANOVA, as implemented in the Fieldtrip function ‘ft_statfun_desamplesFunivariate.m’, and cluster-based Monte Carlo permutation testing with 1000 randomizations.

For source reconstruction, we used a beamformer technique as implemented in the Fieldtrip toolbox (Oostenveld et al., 2010). As a first step we created a head model which is used to estimate the electric field measured by the EEG electrodes. To this end, we segmented a template provided by the Fieldtrip toolbox and estimated a boundary triangle mesh for the volume conduction model. Next, we used a dipoli Boundary Element Method to discretize the brain volume into a grid. For each grid point, we calculated a lead field matrix which was then used to calculate a spatial filter. Following the pre-processing steps described above, the signals were re-referenced to a common-average reference and the spectrum was calculated around stimulus onset (0 ms), separately for pre-stimulus period (−300 −0 ms) and post-stimulus period (0 – 300 ms), with a center frequency of 10 Hz, spectral smoothing factor of 3 Hz and a hanning taper. Then, we computed an inverse filter across all conditions and blocks and applied this filter, during source reconstruction with dynamic imaging of coherent sources (DICS) algorithm, for each condition and block separately. We specified power differences similarly to the electrode-space power analysis described above and performed identical statistical tests (1-way rmANOVA for TP1-TP5), only that clusters here represent grid points. To plot the source analysis results we interpolated the t-values to a template structural MRI. The anatomical labels of the sources were determined using an MNI-based AAL atlas.

To study synchrony between sources, we computed inter-regional spectral coherence by normalizing the magnitude of summed cross-spectral density between two signals and their respective power. Note that DICS was specifically formulated to localize sources coherent with a specific signal and this can be achieved by estimating the cross-spectral density between all EEG channels. Therefore, to compute the coherence with a specific source, we require to specify a channel as reference. Note that it is also possible to specify a specific dipole as reference but depending on the specific voxel chosen as a dipole, the results vary quite substantially. We therefore opted for using channels as reference for the coherence analysis. Here as well, cross-spectral density matrices were calculated for a pre-stimulus period (−300 – 0 ms) and a post-stimulus period (0 – 300 ms) using a center frequency of 10 Hz, spectral smoothing factor of 3Hz and a hanning taper. Next, the coherence differences (SEQ-RND) at each time point (TP1-TP5) were submitted to the same 1-way rmANOVA, to evaluate learning-related changes to α coherence.

### 1.2. Experiment 2

#### 1.2.1. Participants

Twenty-five healthy participants (mean age: 24.8 years, range 20-31; 11 males) took part in the second experiment after giving informed consent. None of the participants had been included in Experiment 1. Participation was financially compensated. All participants were classified as right-handed by means of the Edinburgh Handedness Inventory (Oldfield, 1971) and had normal or corrected to normal vision with no color deficiency. Participants were non-smokers and did not suffer from any mental or neurologic disorder (by self-report). We excluded participants who regularly played an instrument or computer games as well as professional type writers. For the EEG data analysis, one participant was excluded due to a technical error in data acquisition resulting in a total sample of 24 participants. The Participants were blinded to the conditions of the task (as presented below). Experiment 2 was approved by the Ethics Committee of University of Leipzig and was performed in accordance with the Declaration of Helsinki.

#### 1.2.2. Experimental design

Each participant completed three sessions at intervals of at least one week between sessions (Fig. 2A). In each session, participants received 10 Hz tACS to either left M1 (lM1), right cerebellum (rCB), or sham, with the stimulation location counterbalanced between participants. In each session, participants performed a modified version of the serial reaction time task (SRTT) with a different 8-element sequence to prevent cross-over effects. Before and after the experimental blocks, we recorded resting-state sessions, which were not further analyzed for the present work. EEG was collected throughout the experiment in each session. During EEG recordings and task performance, participants were seated comfortably in front of a 17” screen, about 74 cm away, on which visual stimuli were presented. The design of SRTT in Experiment 2, described in Fig. 2C, was similar to Experiment 1. There are the two differences between both experiments: (1) The response-stimulus interval was shorter with 300 ms. (2) The number of SEQ and RND blocks was different (see Fig. 2C). Prior to stimulation, two RND blocks (80 trials each) were practiced. During tACS, we introduced three smaller RND blocks with 40 trials each. As EEG data during tACS was not analyzed, these blocks were used as a behavioral marker for learning during tACS. At stimulation onset, a SMP block (40 trials) was performed, followed by the first smaller RND block. Then four blocks of SEQ with 120 trials each (15 x 8-element sequence) were performed. Here the second RND block was introduced, followed by three blocks of SEQ, the third RND block and a last block of SEQ. Once the stimulation finished, participants performed two large RND blocks (80 trials each) and two SEQ blocks alternately (see Fig. 2C). Next and after a short (~8 min) break, participants performed the completion task which was identical to Experiment 1, except that here we did not ask whether they are sure of their response.

**Figure 2.**
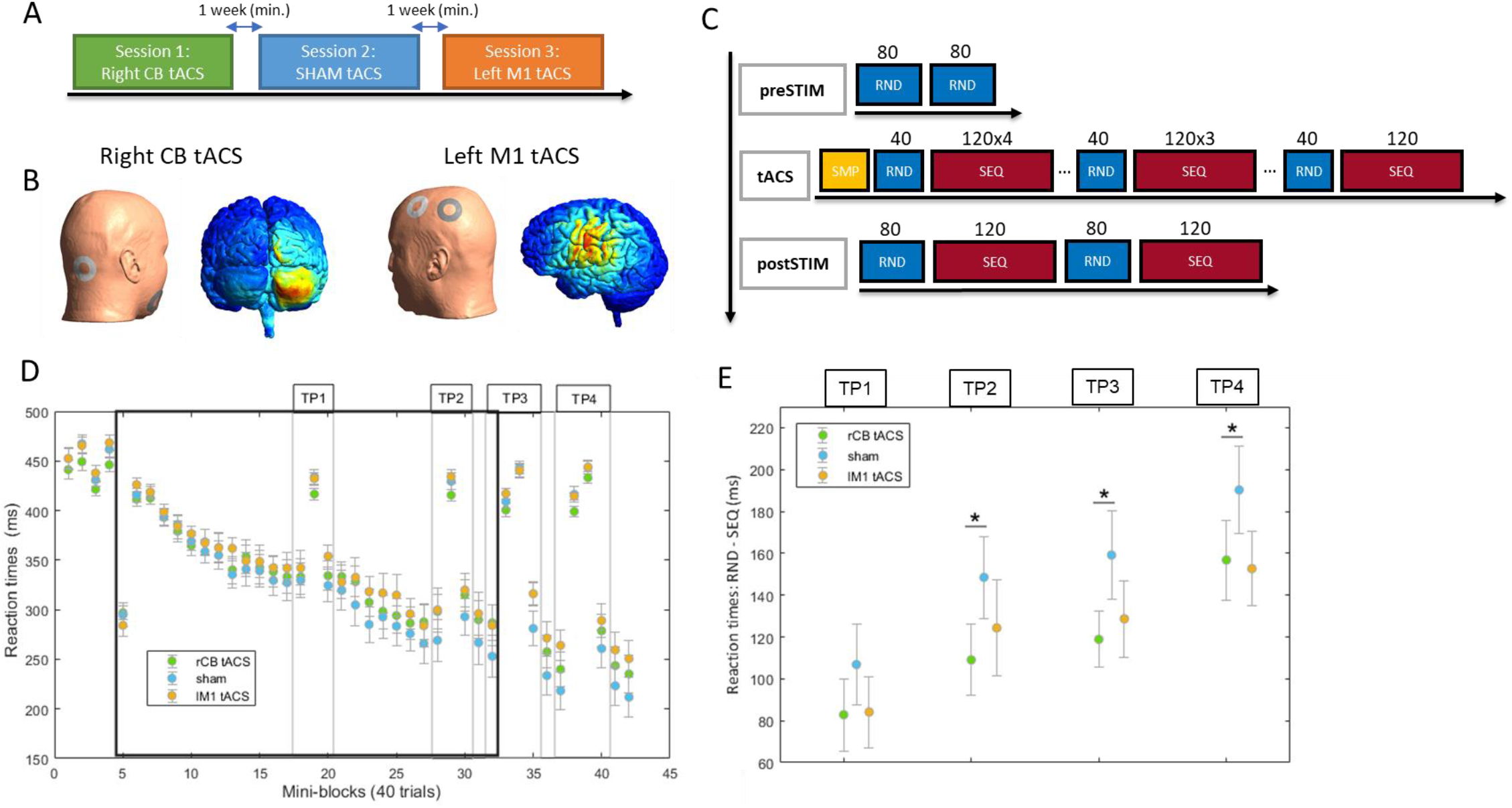
Experiment 2: Experimental design and behavioural results. **A** Experimental sessions. **B** computational modelling of electric field distribution in left M1 and right cerebellum during tACS. **C** Design of serial reaction time task in each session. EEG was recorded throughout the experiment. **D** Reaction times (RT) in mini-blocks averaged across 40 trials each, for each of the stimulation protocols. The duration of tACS is marked with a black frame. Time points (TP) for RT analysis are marked with a light grey frame. Error bars are standard error of the mean (SEM) across subjects. **E** RT differences between RND and SEQ representing better learning performance across TP. Error bars are SEM across subjects. Significant differences are marked with a star (p < 0.05).

#### 1.2.3. Behavioral analysis

We computed reaction times (RTs) and error-rates (ERs) for each of the experimental conditions. Both wrong button-presses and missing responses were regarded as Errors. We excluded trials in which the subject made an error as well as trials in which RTs were bigger and/or smaller than 2.7 SD (corresponding to p < 0.01) of their total average RT. RTs were averaged across mini-blocks of 40 trials, corresponding to five repetitions of the 8-element sequence. We defined four time points (TP) in which sequence learning was assessed by an interruption of a RND block. A learning index was calculated for each TP as follows: the average RT of a RND block minus mean RTs of two SEQ blocks around this RND block (see Fig. 2D). Note that this approach is different compared to Experiment 1, as here only few RND blocks were available. We assessed learning separately during tACS (TP1 and TP2) and following tACS (TP3 and TP4). We then entered these values into 3 x 2 rmANOVA with factors STIM (lM1 tACS, sham; rCB tACS, sham) and factor TIME (TP1 and TP2; TP3 and TP4). The error-rate of each task blocks (SEQ and RND) was computed as the number of errors divided by the number of trials in each block.

#### 1.2.4. Computational modelling of M1 and cerebellar-stimulation locations

We used the SimNIBS software package (http://simnibs.org/), version 2.1 (Saturnino et al., 2019), to simulate the electrode montages to target left M1 and right cerebellum. The head model, provided by the software package, was created using finite element modeling on T1- and T2-weighted MRI images of an exemplary subject resulting in a high-resolution tetrahedral head mesh model containing 8 tissue types. We set the conductivities of brain white matter (WM), gray matter (GM) and of cerebrospinal fluid (CSF) to 0.126 S/m, 0.275 S/m and 1.654 S/m respectively (Opitz et al., 2015). Values of 0.025 S/m and 0.008 S/m were used for the spongy and compact bone of the skull. All tissues were treated as isotropic. The electrical field 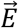 was determined by taking the numerical gradient of the electric potential. For both montages we used ring-shaped electrodes with 5 cm outer diameter, 2.5 cm inner diameter and 3 mm thickness. The total current injected was 1 mA. For lM1 tACS montage, electrodes were placed at EEG locations FC3 and CP3 (Fig. 3B, right montage). For rCB tACS montage, one electrode was placed 1 cm below and 3 cm right to the inion and the other over right mandibula (Fig. 3B, left montage).

**Figure 3.**
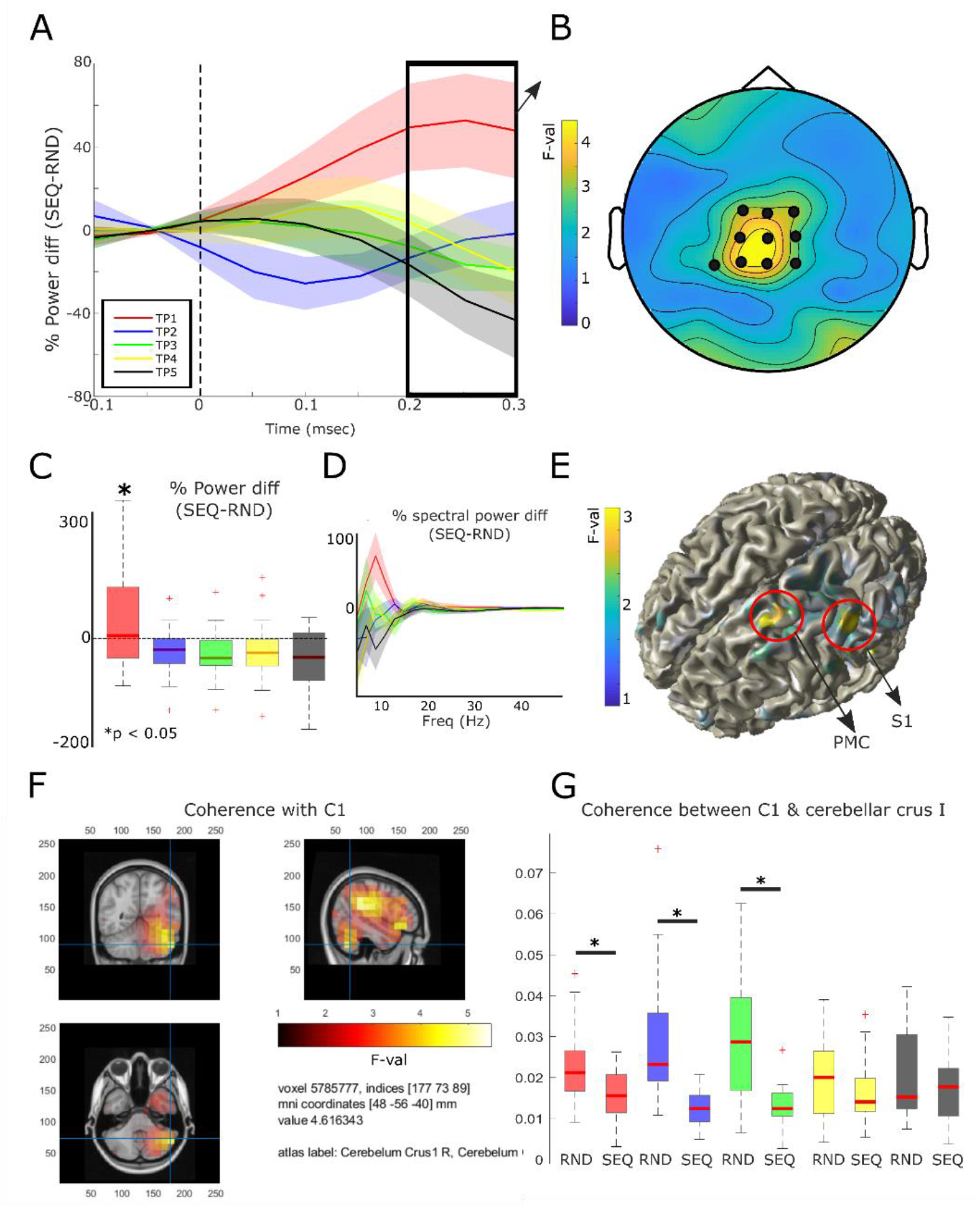
Experiment 1: learning related alpha power modulation and coherence. **A** Time resolved percentage power differences between SEQ and RND blocks for each time point (TP1-TP5). Lines are the mean across subjects and the shade is standard error of the mean (SEM). Significant effects shown in A were found 200-300 ms (black frame) after stimulus onset (dashed line). **B** Topographic map of rmANOVA analysis across TP1-TP5. Electrodes marked with black circle show significant changes with learning. **C** Boxplots for the power differences (SEQ-RND) across TP1-TP5 (same colour code as in A), averaged across 200-300 ms. Stars mark significant differences from 0 (p < 0.05). **D** Percentage power differences across different frequencies between SEQ and RND blocks for each time point (TP1-TP5, same colour code as in A) in the cluster shown B. Values were averaged across 200-300 ms. **E** Source reconstructed activity corresponding to the effects shown in B. **F** Regions showing coherence changes across TP1-TP5 with electrode C1 as reference. **G** Coherence effects across TP1-TP5 (same colour code as in A). Significant differences between conditions are marked with a star (p < 0.005).

#### 1.2.5. Transcranial alternating current stimulation (tACS) protocol

Transcranial alternating current stimulation (tACS) was applied via two ring-shaped conductive rubber electrodes (outer diameter 48 mm, inner diameter 24 mm, area: 15 cm^2^, DC-Stimulator PLUS, NeuroConn, Ilmenau, Germany) with an intensity of 1 mA at 10 Hz (peak-to-peak-amplitude; sinusoidal waveform; 0.07 mA/cm^2^ current density). Ring-shaped stimulation electrodes were used to allow placement around EEG recording electrodes (see below). Prior to electrode placement, the skin surface was treated with high-chloride abrasive electrolyte gel for lowering skin impedance. Impedances were kept below 5 kΩ. Stimulation electrodes were placed either to stimulate rCB or lM1. For rCB tACS, one electrode was placed on the right mandibula and the other 1 cm below and 3 cm right to the inion. For lM1 tACS one ring-shaped electrode was placed around electrode FC3 and one around CP3 rendering the current flow as precisely as possible to C3. For sham, the current was ramped up for 30 s, then stayed at 1 mA for 10 s and ramped down for another 30 s, in order to effectively blind the participants to the experiment protocol. Here, stimulation electrodes were either placed over M1 or cerebellum in a pseudorandomized order across subjects. We used a questionnaire to estimate subjects’ personal assessment of whether they received real stimulation or sham.

#### 1.2.6. EEG recordings and pre-processing

EEG was recorded using Ag/AgCl electrodes embedded in a 64-channel cap and connected to an eego^TM^ amplifier (ANT Neuro b.v., Hengelo, the Netherlands) with a sampling rate of 512 Hz. Eye movements were recorded with an electrooculogram (EOG) below the left eye. Electrodes were placed according to an extension of the international 10–20 system. The EEG was recorded against an online reference electrode in CPz. All electrode impedances were kept below 5 kΩ.

Pre-processing and all subsequent analyses were performed using in-house Matlab (The Mathworks®, Natick, MA) scripts and the EEGLAB toolbox. EEG signals were analyzed before and after tACS but not during stimulation due to artifacts (Noury et al., 2016). We first applied a high pass filter (lower edge of the frequency pass band 1 Hz, higher edge of the frequency band 49 Hz, FIR Filter order = 5000) to remove slow drifts and power line noise. Signals were then re-referenced offline to the average of the signal from left and right mastoids and the signal from electrode CPz was re-calculated. Next the signals were segmented into 3 s epochs (−1 s to 2 s around stimulus onset). Based on ICA, we visually identified 3-4 components related to eye blink artifacts and removed them. Additional artifacts were removed using a simple threshold (−70 μV, +70 μV) on the filtered data.

#### 1.2.7. EEG spectral power and source analysis

We computed the power spectrum of the EEG signals using the Morlet wavelet as implemented in Fieldtrip, in the exact same procedure as in Experiment 1. Here we focused our analysis on EEG collected after tACS to assess the influence of stimulation on learning. To this end, we computed power differences between SEQ and RND trials in TP3 and TP4 (TP1 and TP2 were during tACS, see Fig. 3D). Statistical analyses were performed using non-parametric cluster-based Monte Carlo permutation testing with 1000 randomizations, comparing power differences between rCB and sham, as well as between lM1 and sham, at each TP. Sources responsible for producing oscillatory activity and inter-regional spectral coherence were identified exactly as in Experiment 1 (see section 1.1.6 above for further details).

## 2. Results

### 2.1. Experiment 1

#### 2.1.1. Behavioral results

One subject was excluded from further analyses due to large error-rates, more than 2.7 SD from the group mean, resulting in a sample of 24 subjects. To assess motor sequence learning we subjected the reaction times (RTs) to repeated measures ANOVA (rmANOVA) with factors COND (SEQ, RND) and TIME (5). We found a main effect of COND (F1,23 = 156.5, p < 0.001) and main effect of TIME (F4,92 = 30.6, p < 0.001) as well as a COND x TIME interaction (F4,92 = 10.2, p < 0.001) suggesting that subjects improved their performance in sequence blocks compared to the random blocks with time (Fig. 1C). As error rates (ERs) were not distributed normally, we were not able to apply rmANOVA as with the RTs. We, therefore, assessed condition differences in each block, taking all mini-blocks into account for calculation of the total amount of errors per block. We found larger error-rates during RND blocks compared to SEQ blocks (see supp. Fig. 1) in TP1 (p = 0.01), TP4 (p = 0.05), with the largest difference at the end of the task at TP5 (p < 0.001). These results reveal clear learning of the motor sequence reflected by faster and more accurate performance in SEQ blocks.

After the SRTT, we examined subjects’ knowledge of the sequence using the completion task. Subjects were tested on 32 completion trials and gave correct and correct assured (a correct response followed by a “yes” answer to the question: “are you sure?”) responses. Note that chance level is 33%. The correct response rate was 67.2% ± 15.9% and correct assured responses was 40.3% ± 26.9%, suggesting that most subjects could successfully reproduce at least some of the sequence.

#### 2.1.2. Learning-related effects on post-stimulus power

We investigated learning related changes in oscillatory power using a time-frequency analysis. Based on our hypotheses outlined above, we examined differences in learning-related power across TIME (TP1-TP5), focusing on α (8-13 Hz) frequency band, 0-300 ms time window following stimulus onset. Note that this time window was chosen since the average RT at the end of the task is around 300 ms (Fig. 1C). To this end, we subjected the power differences (SEQ-RND) in TP1-TP5 to a 1-way rmANOVA, as implemented in the Fieldtrip function ‘ft_statfun_desamplesFunivariate.m’ and cluster-based permutations. We found a significant effect in a central cluster (Fig. 3B, 200 – 300 ms, cluster level p = 0.04), which reflected changes in learning-related (SEQ-RND) α power across TP1-TP5 (Fig. 3A). Specifically, α power in this cluster increased in TP1 (t21 = 2.3, p = 0.03) and tended to decrease in TP5 (t21 = −2.1, p = 0.05, Fig. 3A, 3C), with no changes observed in TP2-TP4 (p > 0.2). In addition, α power significantly decreased in SEQ5 compared to SEQ1 (diff: 41.5, t21 = 3.2, p = 0.004). Importantly, we tested whether this effect was specific to α, and found no differences (all p > 0.1) in other frequency bands (θ: 4-8 Hz, β: 13-30 Hz, γ: 30-50 Hz) (Fig. 3D). These results suggest that motor memory encoding is characterized by an early α power increase as well as late α power decrease. There were no significant correlations between learning-related α power changes and behavioral markers of learning indexed by RT differences (all p > 0.09).

Using the beamformer technique, we reconstructed sources of learning-related α power across time (TP1-TP5) using the same rmANOVA at the electrode-level described above. Activity was evident in left PMC (peak voxel at F4,21 = 3.6, see Fig. 3E), somatosensory cortex (S1, peak voxel at F4,21 = 3.6, see Fig. 3E) and temporoparietal junction (TPJ, peak voxel at F4,21 = 3.6, not shown). We also tested whether learning-related oscillatory α power changes were evident after the motor response and before the next stimulus appears. Here, we had a constant 500 ms response-stimulus interval (see methods above). Using the same statistical analysis as with the post-stimulus α power above, we found no significant clusters showing learning-related α power across time (TP1-TP5).

#### 2.1.3. Connectivity between neural sources

Next, we computed the coherence between neural sources. We specified electrode C1 as reference (see details in section 1.1.6), to account for both left PMC and S1 sources. We then asked whether learning-related (SEQ-RND) α coherence changes as learning progresses, similar to learning-related α power. To this end, we submitted the coherence differences (SEQ-RND) at each time point (TP1-TP5) to the same rmANOVA analysis above. We found a large cluster in right TPJ, extending to inferior parts of the peri-central gyrus (cluster level p < 0.05, peak voxel at F4,21 = 5.4), inferior frontal gyrus (IFG, F4,21 = 4.5), as well as right posterior cerebellum (F4,21 = 4.6, Fig. 3F). Post-hoc t-tests show that coherence in this cluster was stronger during RND compared to SEQ in TP1-TP3 (Fig. 3G, all p < 0.005), while later in learning (TP4 and TP5) no differences in coherence between RND and SEQ were evident (p > 0.1). These results suggest that connectivity between PMC, S1 and cerebellum is reduced during early SEQ blocks.

### 2.2. Experiment 2

None of the subjects reported any adverse effects during or after stimulation. We asked subjects whether they experienced phosphenes, which would be a sign of visual cortex stimulation (Kar and Krekelberg, 2012). Seven out of 25 subjects reported phosphenes during either lM1 tACS (N = 2) or rCB tACS (N = 5). Three of the seven claimed to see phosphenes during sham. There were no reports on pain or dizziness due to stimulation. During real tACS when asked whether the session was sham, subjects correctly answered “no” in 48% of all real tACS sessions, which shows that subjects were blinded to the intervention. As subjects performed the task a total of three times, we also checked whether the order of the sessions affected learning by subjecting reaction time (RT) differences between two SEQ blocks and a “sandwiched” RND block, at each time point (TP), to a repeated measures ANOVA (rmANOVA) with factors SES (sessions 1-3), and TIME (TP1-TP4). There was no main effect of SES (p = 0.18) or SES x TIME interaction (p = 0.09), suggesting no effect of session order on motor sequence learning in this experiment. A test for the effect of sequence complexity on learning speed is reported in the supplementary materials.

#### 2.2.1. Right cerebellar tACS leads to diminished learning

To assess performance changes in the motor sequence learning task due to rCB and lM1 tACS, we first subjected the RT differences between two SEQ blocks and a “sandwiched” RND block, at each time point (TP), to a rmANOVA analysis with factors STIM (rCB tACS, lM1 tACS and sham), and TIME (during stimulation: TP1 and TP2; following stimulation: TP3 and TP4). Both during and following tACS we found a main effect of TIME (both comparisons: F1,24 > 20.9, p < 0.001), reflecting a significant increase from TP1 to TP2 and from TP3 to TP4 (see Fig.3E). This suggests that subjects learned the underlying sequence across all stimulation protocols. No effects of STIM or STIM x TIME interactions were evident (p > 0.1).

We then asked if each of the stimulation protocols differed from sham. To this end we employed separate rmANOVA analyses for rCB tACS vs. sham and for lM1 tACS vs. sham as factor STIM and a factor TIME similar to the analysis above. Here as well we found a main effect of TIME for TP1 and TP2 (both comparisons: F1,24 > 13.1, p < 0.002) as well as for TP3 and TP4 (both comparisons: F1,24 > 21.4, p < 0.0002). Importantly, a main effect of STIM was evident only for the comparison between rCB tACS and sham, both during stimulation (TP1 and TP2: F1,24 = 6.2, p = 0.02), and following stimulation (TP3 and TP4: F1,24 = 5.9, p = 0.02). No main effect of STIM was evident for the comparison between lM1 tACS and sham (p > 0.1).

To explore when rCB tACS affects learning, we compared between rCB tACS and sham in each TP separately. Note that no correction for multiple comparisons was applied to these post-hoc tests. We found reduced learning comparing rCB tACS to sham, expressed by a lower RND-SEQ RT difference in rCB tACS (see Fig. 3E), in TP2 (t24 = 2.6, p = 0.015), TP3 (t24 = 2.3, p = 0.03) and TP4 (t24 = 2.1, p = 0.04). We also explored differences between stimulation sites (rCB tACS vs. lM1 tACS) but no effects were found (p > 0.7). There were no STIM x TIME interactions (all p > 0.3) suggesting that improved learning with time was independent from stimulation.

We further explored whether tACS affected general task performance during RND blocks. Here we found that in TP1 (p = 0.08), and TP4 (p = 0.06), RTs tended to be faster during rCB tACS when compared to sham (see Fig. 3D). No such effects were evident for lM1 tACS (p > 0.6). Together, these results suggest that while rCB tACS leads to reduced sequence learning, it tends to also speed up general task performance.

Next, we analyzed the effect of rCB and lM1 tACS on error-rates. Subjects made significantly more errors during RND (median ± std: 3.2% ± 1.8%) compared to SEQ blocks (0.14 ± 0.29%; Wilcoxon signrank: p < 0.0001) across all stimulation protocols, indicating successful learning of the sequence. The error-rates did not distribute normally which prevented an analysis with the same rmANOVA as the RTs. To this end, we performed Wilcoxon signrank tests in each TP in the RND blocks only. No difference between either rCB or lM1 tACS and sham were observed (all p > 0.1). Performance in the completion task was high across stimulation protocols with median percentage correct responses for rCB tACS: 83.3%, lM1 tACS: 87.5% and sham: 87.5%. No differences in completion task performance between lM1 or rCB tACS and sham were evident (Wilcoxon signrank test: all p > 0.1).

#### 2.2.2. Alpha entrainment following 10Hz tACS

We excluded one subject from all EEG analyses described below due to extreme outliers (more than 3 SD) in α power across conditions, resulting in a total of 24 subjects. In accordance with previous reports (Zaehle et al., 2010), we expected that 10Hz tACS would entrain local oscillatory activity, reflected by increased α power in experimental blocks directly following the stimulation. To this end, we first computed α power differences between preSTIM RND blocks and postSTIM RND blocks. We then compared these differences between stimulation protocols. No significant clusters were found. As these changes might be affected by both tACS and learning, we directly compared α power differences between stimulation protocols (rCB tACS vs. sham, lM1 tACS vs. sham), during the first post-stimulation RND block. A significant α power increase in electrodes O2 and Oz (p = 0.006, Fig. 4A-B) was evident following rCB tACS compared to sham. This effect was not specific to α, as significant power differences were also observed in the θ frequency band (4-8 Hz) in these electrodes, expanding to electrodes O1 and O3 as well as to a left centro-parietal cluster (see supp. Fig 2.). No effects were found in β (13-30 Hz) or γ (30-48 Hz) frequency bands (all p > 0.1). No differences were found following lM1 tACS compared to sham. We then used beamforming to locate α power changes between stimulation protocols. Comparing rCB tACS to sham, we found a cluster in left inferior parietal cortex (peak voxel at t23 = 5.2, cluster level p = 0.02, family-wise error corrected). No sources were found for lM1 tACS compared to sham.

**Figure 4.**
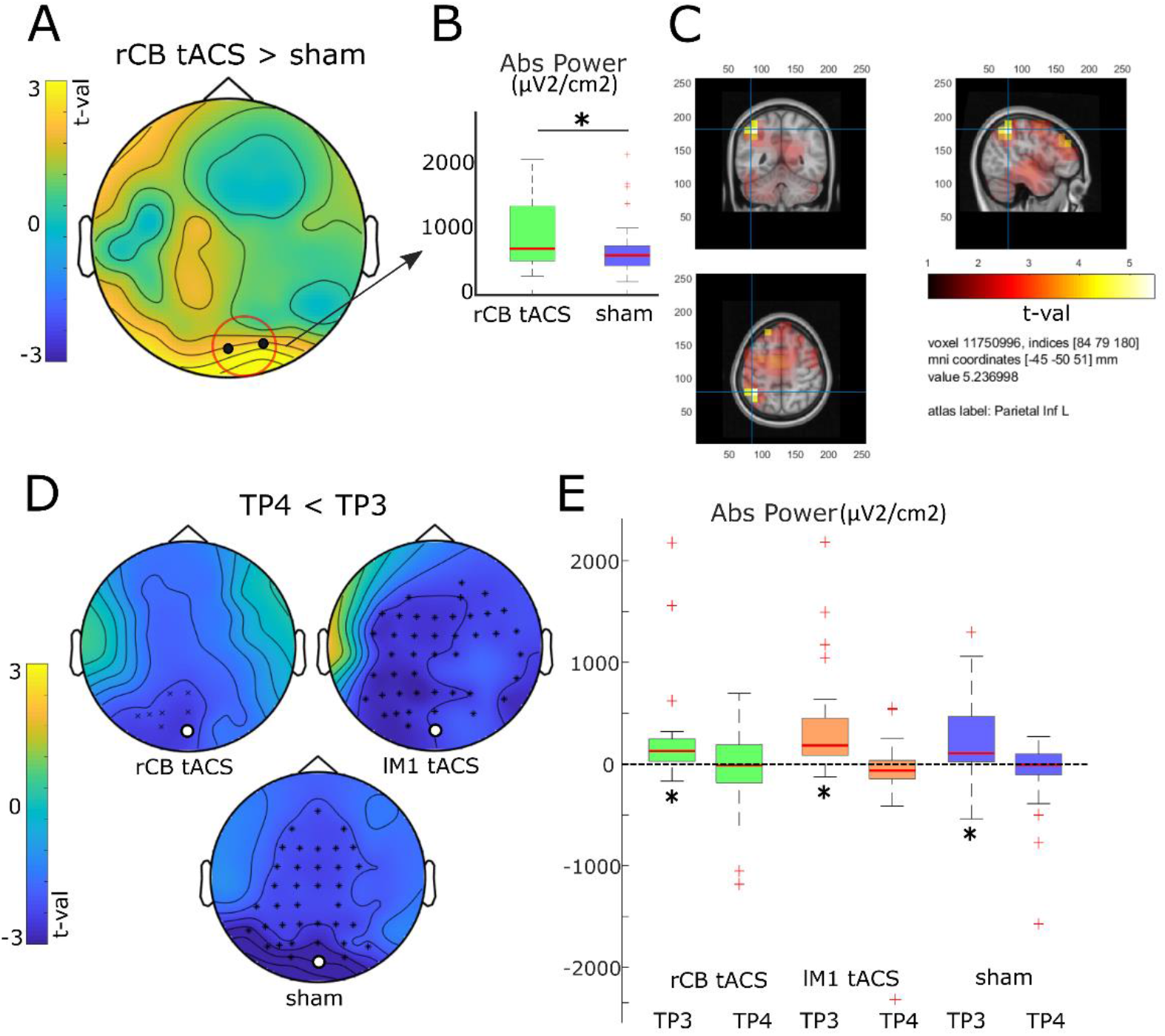
Experiment 2: Alpha power modulation following 10 Hz tACS. **A** Topographic map for alpha power differences between rCB tACS and sham in the first RND block following stimulation. Significant electrodes are shown in black and marked with a red circle. **B** Boxplots for the absolute power (μV^2^/cm^2^) averaged across electrodes O2 and Oz (shown in A). The star marks a significant difference (p = 0.006). **C** Source reconstruction of power differences between rCB tACS and sham, corresponding to the map in A. **D** Topographic maps for each stimulation protocol showing decreased alpha power from TP3 to TP4. **E** Boxplots for the absolute power (μV^2^/cm^2^) across the different stimulation protocols and time points in an example electrode Oz (marked in D with a white circle).

#### 2.2.3. Effects of lM1 and rCB tACS on learning-related α power

Next, we examined in the post-stimulation task blocks, whether tACS affected learning-related α power. We focused on a time window of 0-200 ms following stimulus onset. Note that here the time-frame is shorter compared to Experiment 1, since participants were much faster towards the end of the task (see Fig. 3D and Fig. 1C). We then computed the power differences (SEQ-RND) in TP3 and TP4. We first asked whether α power changes are evident when learning progresses, as we found in the first experiment. Comparing TP3 to TP4 in each of the stimulation protocols, we found a broad α power decrease following lM1 tACS and sham (Fig. 4D-E), and in fewer electrodes following rCB tACS. This effect was reflecting learning-related increase in TP3 (e.g. electrode Oz; rCB tACS: t23 = 2.6, p = 0.02, lM1 tACS: t23 = 3.3, p = 0.004, sham: t23 = 3.0, p = 0.007) which vanished at TP4 (all p > 0.2). Notably, the early α power increase replicates the results of Experiment 1.

Then we asked whether tACS affected learning-related α power, by comparing lM1 tACS and rCB tACS to sham. We found a significant increase in learning-related (SEQ-RND) α power in a left fronto-central cluster (cluster level p = 0.04), when comparing rCB tACS to sham in TP4 (Fig. 5A-B). No effects were found in θ, β or γ frequency bands (all p > 0.1, see also supp Fig. 3 for learning-related power in this cluster across different frequencies). Source reconstruction revealed that this effect in TP4 originated from a cluster in left PMC (peak voxel t = 3.1, Fig. 5C). No such differences were observed between lM1 tACS and sham. No significant differences in learning-related α power between stimulation protocols were found in TP3.

**Figure 5.**
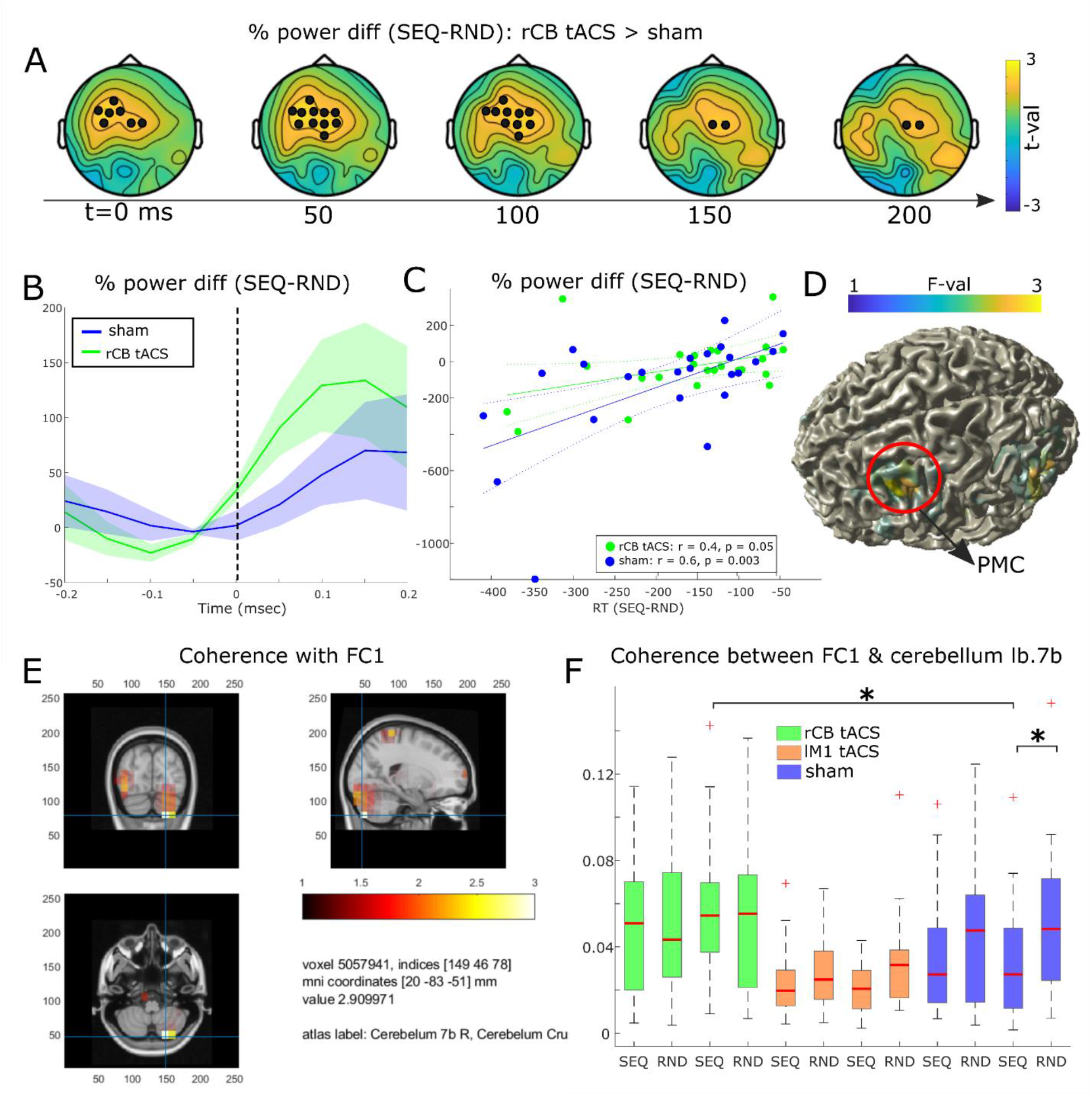
Experiment 2: The effect of rCB tACS on learning-related alpha power. **A** Topographic maps for learning-related (SEQ-RND) alpha power differences between rCB tACS and sham, in a 200 ms time frame following stimulus onset (t = 0 ms). Significant electrodes are marked with a black circle. **B** Time resolved power differences between SEQ and RND blocks in rCB tACS and sham averaged across the cluster shown in A (t = 100 ms). Lines are the mean across subjects and the shade is the standard error of the mean. Stimulus onset is marked with a dashed line. **C** Correlation between a learning measures (RT differences) and power differences (SEQ-RND), following rCB tACS (green) and sham (blue), in TP4. **D** Source reconstruction corresponding to the maps in A. **E** Cerebellar cluster showing coherence effects with the seed in left PMC, shown in D. **F** Boxplots for coherence effects between left PMC and right cerebellum. Significant differences are marked with a star.

As learning effects may be evident even prior to stimulus presentation, we also tested whether tACS affected learning-related α power before stimulus onset (−200-0ms). We found that α power in electrodes F5 and FC5 was stronger (p = 0.02 and p = 0.03 resp., data not shown) following rCB tACS compared to sham, suggesting that rCB tACS effect on learning-related α power is not specific for the period prior to the motor response but is probably a general effect.

To investigate how changes in α power may relate to individual performance, we correlated RT differences in TP4 with learning-related α power post-stimulus (no baseline correction was applied) in the fronto-central cluster. We found that learning-related α power had a tendency for a positive correlation with individual performance (r = 0.4, p < 0.05), following rCB tACS (see Fig. 5E). Similarly, a significant positive correlation was evident (r = 0.6, p = 0.003) following sham (Fig. 5E). This means that in both rCB tACS and sham, stronger α power decrease during learning was associated with better learning (larger RT differences). No correlations were found following lM1 tACS.

#### 2.2.4. Effects of rCB tACS on premotor-cerebellar connectivity

Next, we computed α coherence between neural sources. Here we specified electrode FC1 as reference to account for the power effects found above. We then compared the differences in coherence between rCB tACS to sham at TP3 and TP4 (both SEQ-RND). We found at TP4, small clusters in right cerebellar lobule VIIb (Fig. 5D, peak voxel at t = 2.9) and left occipital cortex (peak voxel at t = 3.4, not shown). Post-hoc t-tests in cerebellar crus I (see Fig. 5F) showed that coherence during SEQ at TP4 was stronger following rCB tACS when compared to sham (t23 = 3.4, p = 0.002), with no such differences in RND (p > 0.6). In addition, following sham, coherence in RND was stronger compared to SEQ (t = 4.3, p < 0.001), replicating the findings of Experiment 1, whereas no differences were evident at TP4 between SEQ and RND following rCB tACS (p > 0.8). Together, these results suggest that rCB tACS leads to abolished differences in connectivity between left PMC and right cerebellum relevant for motor learning, by enhancing connectivity during SEQ blocks.

## Discussion

In this study we demonstrated a strong link between alpha oscillations (α, 8-13 Hz) in the premotor-cerebellar loop and motor sequence learning. We showed that α power over left premotor cortex (PMC) and sensorimotor cortex (SM1) increased early-on and decreased as learning progressed. At the same time, α coherence between left PMC, S1 and right cerebellar crus I was weaker in sequence (SEQ) compared to random (RND) blocks in early learning blocks. This could indicate a functional decoupling within a cerebellar-motor loop guided by α oscillations and underlying motor learning. In addition, 10Hz transcranial alternating current stimulation (tACS) to right cerebellum (rCB) during sequence learning interfered with learning and led to a specific learning-related α power increase over left PMC. Importantly, condition differences in α coherence between left PMC and right cerebellar lobule VIIb evident following sham, were abolished following rCB tACS, through a specific increase during SEQ blocks. Together these results suggest that: (1) α oscillations in the cerebellar-motor loop are modulated during motor sequence learning (2) external entrainment of cerebellar α oscillations interferes with motor learning, through increased α coherence in premotor-cerebellar connections and increased α power in left PMC.

### Learning modulates α power in premotor- and sensorimotor cortex

In Experiment 1, we found that learning-related α power in left PMC and S1 increased early-on and decreased as learning progressed. Importantly, this effect was found prior to the response with no apparent changes to α power in a similar period pre-stimulus. This corresponds to previous findings showing that the more a movement is performed automatically, the less α power desynchronizes over motor areas (Boenstrup et al., 2014). Rueda-Delgado et al. (2019) have recently shown that following bimanual coordination task, α power in resting-state EEG was also decreased in sensorimotor and premotor areas. Based on the functional inhibition theory by Jensen and Mazaheri (2010), postulating that decrease in α oscillations reflects release of inhibition of task-relevant areas, it is plausible that α power decrease represents activation of PMC during encoding of the motor sequence. Indeed, previous fMRI studies of motor sequence learning consistently show increased activity in PMC as learning progresses (Bapi et al., 2006; Grafton et al., 2002; Orban et al., 2010).

We further found that α coherence between left PMC, right TPJ and cerebellar crus I was significantly weaker during SEQ compared to RND. Previously, we showed that reduced α phase coupling in SEQ compared to RND, interpreted to reflect global functional decoupling, is important for encoding the sequence into memory (Tzvi et al., 2018). We therefore suggest that α decoupling between left PMC, right TPJ and cerebellar crus I serves to integrate motor representations in motor cortical areas, attentional processes in TPJ (Corbetta and Shulman, 2002), and internal model representations in the cerebellum (Ito, 2006) during motor learning.

### Entrainment of posterior α oscillations following 10 Hz tACS to right cerebellum

We found increased α power in occipital electrodes, in RND blocks directly following rCB tACS compared to sham. The source of this effect originating in left inferior parietal lobe (IPL), suggests that entrainment of oscillations by rCB tACS elicited a network effect. The IPL is important for sensory processing and sensorimotor integration and is directly disynaptically connected to the cerebellum (Clower et al., 2001). Thus, in accordance with the behavioral results showing a tendency for improved general task performance following rCB tACS, it is plausible that entrainment of cerebellar α oscillations resulted in changes to IPL-cerebellar loop underlying visuomotor processes. This speculation should be further investigated in future studies focusing on modulation of stimulus-response processes. Interestingly, in a group of Parkinson’s and essential tremor patients, alpha tACS to the cerebellum led to a significant entrainment of tremor frequency (Brittain et al., 2015). Given that rCB tACS was applied superficially, it is likely that the cerebellar cortex is the most affected by the stimulation. Purkinje cells, as the sole output of the cortex, were strongly entrained by AC stimulation in a large range of frequencies, shown by extracellular recordings from vermis lobule 7 in anesthetized rats (Asan et al. 2020).

### The effect of 10 Hz tACS on motor learning performance

The results of Experiment 1 suggested that α entrainment in left PMC and SM1, as well as coherent regions, right TPJ and cerebellar crus I, through external stimulation would disrupt learning. Indeed, in Experiment 2 we found that 10Hz rCB tACS, but not lM1 tACS, led to declined learning performance when compared to sham. Only few studies investigated the effect of cerebellar tACS on motor performance (Miyaguchi et al., 2018; Naro et al., 2016). These studies show that higher frequency tACS to the cerebellum led to improved motor performance, but contrary to our results, found no effect at 10Hz. Notably, 10Hz CB tACS was applied at rest, whereas we stimulated during motor learning which is likely to critically impact the results (Ruhnau et al., 2016).

### The effect of 10 Hz tACS on learning-related α oscillations

Comparing between rCB tACS to sham, we found a relative increased learning-related (SEQ-RND) α power over fronto-central electrodes following rCB tACS. Correlation between this effect and behavioural measures of learning revealed that better learners had a stronger α decrease, following both sham and rCB tACS (but not lM1 tACS). This result further corroborates the findings of Experiment 1, linking α power decrease with encoding of the motor sequence. Source reconstruction of the effect over fronto-central electrodes revealed a left PMC cluster, similar to the results of Experiment 1. The relative learning-related power increase following rCB tACS could indicate reduced activity in PMC resulting in impaired learning.

Importantly, we found that learning-specific coherence between left PMC and right cerebellum is stronger following rCB tACS compared to sham. This suggests that entrainment of cerebellar oscillations by tACS led to increased coherence in the premotor-cerebellar loop during learning, ultimately leading to increased α power at PMC. Projections from cerebellum to premotor cortex, shown in macaques using retrograde transsynaptic transport of a neurotropic virus (Hashimoto et al., 2010) suggest that cerebellum strongly impacts higher cognitive functions represented in PMC. In addition, we previously found using fMRI, negative modulation of interactions from motor and premotor cortex to cerebellum by motor sequence learning (Tzvi et al., 2015, 2014), as well as specific interaction from right cerebellum to left PMC, leading to increased left PMC activity, associated with learning in a visuomotor adaptation task (Tzvi et al., 2020). We thus speculate that α mediated communication in the PMC-cerebellar loop serves to integrate perceptual and motor components for motor learning. Taking the results of both experiments together, we propose the following mechanism: entrainment of cerebellar α oscillations by 10Hz cerebellar tACS leads to increased cerebellar output activity, which results in enhanced α coherence with PMC. The resulting α power increase in PMC, supposedly reflecting PMC inhibition is linked to the observed diminished sequence learning.

### Limitations

The computational model suggests that the montage we used for cerebellar tACS elicits a response mostly in the cerebellum. Possibly, such a montage may also elicit a response in the near-by visual cortex. However, we believe that this not the case in Experiment 2 since only very few subjects experienced phosphenes only under cerebellar tACS. For technical reasons it is impossible to know if local cerebellar oscillations were entrained by rCB tACS. We showed, however, significant entrainment of occipital electrodes, also predicted by the computational model (Fig. 3B), which were closest to the stimulation electrode targeting the cerebellum. This suggests that electrodes placed inferior to occipital electrodes might be able to detect cerebellar oscillations (Todd et al., 2018). In addition, entrainment of other cortical regions, especially left IPL, which is tightly connected to right cerebellum (Bostan et al., 2013), as well as the observed coherence between PMC and cerebellum suggests that tACS targeting the cerebellum could entrain local oscillatory activity. Surprisingly, 10Hz tACS to left M1 did not significantly affect learning compared to sham as expected from previous work (Fresnoza et al., 2020; Pollok et al., 2015). In previous studies, stimulation electrodes were placed over left M1 (at C3 or lM1 measured by TMS) and right orbita possibly leading to a distributed effect over not just M1 but PMC and large portions of prefrontal areas (e.g. Opitz et al., 2015), which might have led to the positive learning effects. However, as we found no evidence for local entrainment of mu oscillations following lM1 tACS, we suspect that placement of stimulation electrodes over FC3 and CP3 might have led the current to be shunted across the scalp, due to relative proximity of the stimulation electrodes resulting in a weaker electric field strength at M1.

### Conclusions

The results of this study suggest that α oscillations mediate interactions in the premotor-cerebellar loop and thus modulate motor sequence learning. Specifically, a gradual decrease of learning-related α power in left premotor cortex, and weaker learning-specific coherence with right cerebellum suggests a functional decoupling of the premotor-cerebellar loop underlying motor learning. When 10Hz tACS was applied to right cerebellum, learning-related α power increased in left premotor cortex and was more coherent with right cerebellum compared to sham. In sum, these results suggest that interactions within a premotor-cerebellar loop, underlying motor sequence learning, can be modulated through external entrainment of cerebellar oscillations leading to both behavioral and physiological changes.

## Acknowledgements

The authors declare no competing financial interests. We would like to thank Mourad Zoubir, Gregor Spitta and Susanne Schellbach for their assistance with EEG data collection in Experiment 1. This work is supported by the German-Israeli foundation, grant G-2510-421.13/2018 to ET and by the DFG grant TZ 85/1-1 to ET.

