## supplementary materials for "The role of alpha oscillations in a premotor-cerebellar loop in modulation of motor learning: insights from transcranial alternating current stimulation"

1. **Experiment 2: No effect of sequence complexity on declined learning following rCB tACS**

We examined whether the specific sequence influenced learning performance by submitting RTs of each sequence to rmANOVA with factors SEQ (SEQ1, SEQ2, SEQ3) and TIME (TP1-TP4). Here we found a main effect of SEQ (F_2,48_ = 5.4, p = 0.007) suggesting that one or more of the sequences were learnt faster. The interaction SEQ x TIME was not significant (p = 0.4) which means that while one of the sequence might have been easier than other, this didn’t affect improvements in learning with time. Post-hoc t-tests revealed that one of the sequences was indeed easier than the others (see supp. Table 1). As sessions and sequences were pseudorandomised and counterbalanced across stimulation protocols (sham, lM1 tACS, rCB tACS) and there was no ceiling effect in any of the sessions, we believe that the easier sequence did not affect the observed declined performance following rCB tACS reported in the main text. In addition, at TP4, differences between the sequences were not evident anymore, suggesting that late learning did not differ between different sequences.

*Supplementary Table 1: Post-hoc t-tests comparing RTs between sequences in each learning point.*

|  | **TP1** | | **TP2** | | **TP3** | | **TP4** | |
| --- | --- | --- | --- | --- | --- | --- | --- | --- |
|  | *t* | *p* | *t* | *p* | *t* | *p* | *t* | *p* |
| **SEQ1, SEQ3** | 2.3 | 0.03 | 2.4 | 0.02 | 2.5 | 0.02 | 1.1 | 0.3 |
| **SEQ2, SEQ3** | 3.2 | 0.004 | 2.4 | 0.02 | 3.2 | 0.004 | 1.9 | 0.07 |
| **SEQ1, SEQ2** | 0.6 | 0.5 | 0.4 | 0.7 | 0.8 | 0.4 | 0.9 | 0.4 |

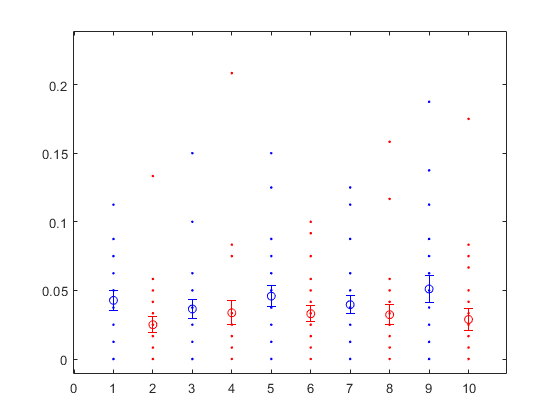

RND SEQ

**Supp. Fig 1.** Error-rates in Experiment 1 averaged across subjects and trials in each time point corresponding to Fig 1D in the main text. Conditions are colour coded: Random-blue. Sequence–red. Error bars are standard errors of the mean.

TP1

TP2

TP3

TP4

TP5

**

*

*

** p < 0.01

* p < 0.05

Error-rate

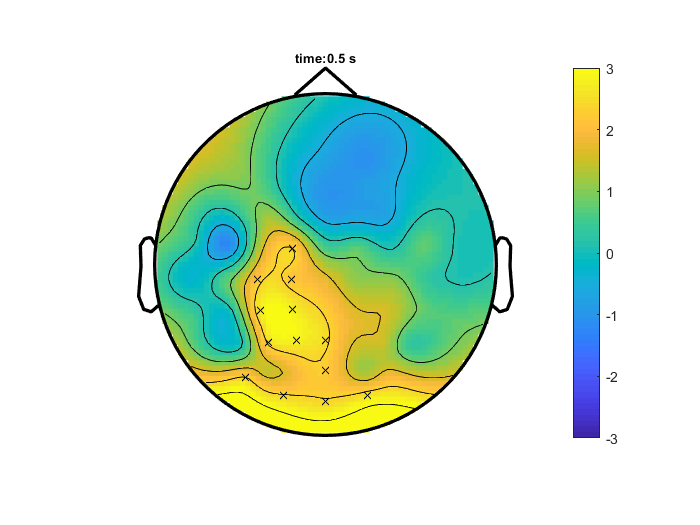

**Supp. Fig 2.** Topographic map for theta power differences between rCB tACS and sham in the first RND block following stimulation. Significant electrodes are marked with a ‘x’.

rCB tACS vs. sham at t = 100 ms

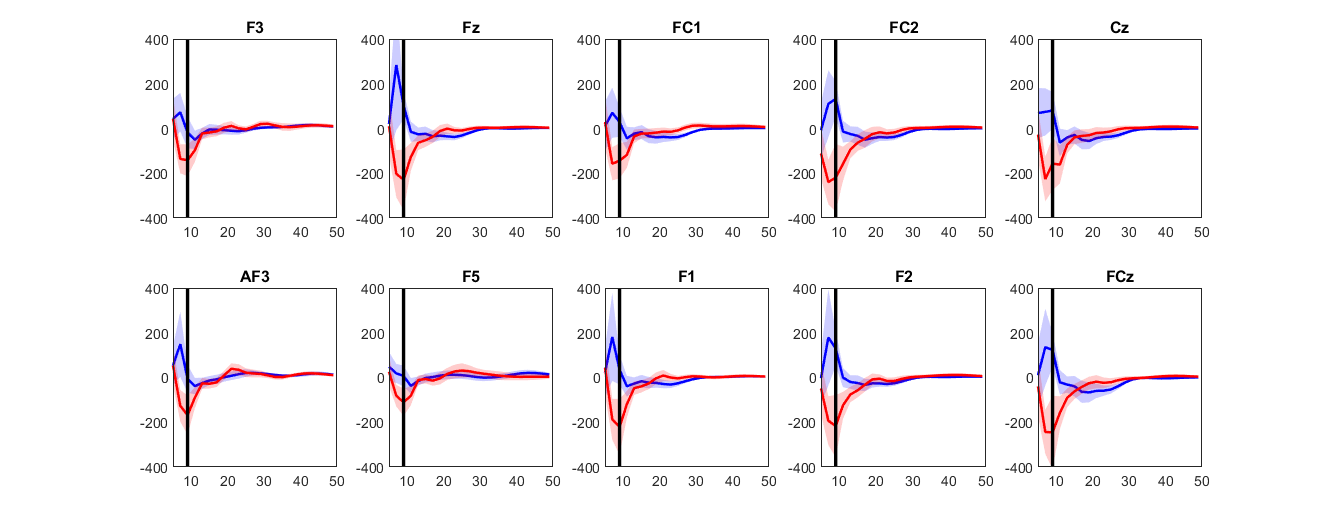

**Supp. Fig 3.** Spectral percentage power differences in TP4, in the electrodes of the cluster shown in Fig. 5A in the main text.

Freq (Hz)

% Power diff (SEQ-RND)
